## Supplemental for "LSD and psilocybin flatten the brain’s energy landscape: insights from receptor-informed network control theory"

[H] Institució Catalana de la Recerca i Estudis Avançats (ICREA), Passeig Lluís Companys 23, Barcelona, Spain

[I] Department of Neuropsychology, Max Planck Institute for Human Cognitive and Brain Sciences, Leipzig, Germany

[J] School of Psychological Sciences, Monash University, Melbourne, Clayton, Australia

[K] Department of Psychiatry, University of Oxford, Oxford, United Kingdom

[L] Center of Music in the Brain (MIB), Clinical Medicine, Aarhus University, Denmark

[M] Centre for Eudaimonia and Human Flourishing, University of Oxford

[N] Department of Radiology, Weill Cornell Medicine, New York, USA

***Corresponding author**

***Assessing the stability of clustering***

As mentioned in the main text, we performed 50 repetitions of *k*-means clustering, choosing the lowest error solution. We wanted to further ensure that this solution was a consistent global minimum by repeating the former process 10 times and comparing the adjusted mutual information[1] (AMI) shared between the 10 partitions. The partition that had the maximum sum of AMI scores with all other 9 partitions was selected for analysis. More importantly, this process confirmed that *k*-means clustering was consistent and arriving at stable solutions since AMI between partitions was generally high (> 0.90; SI Figure 1).

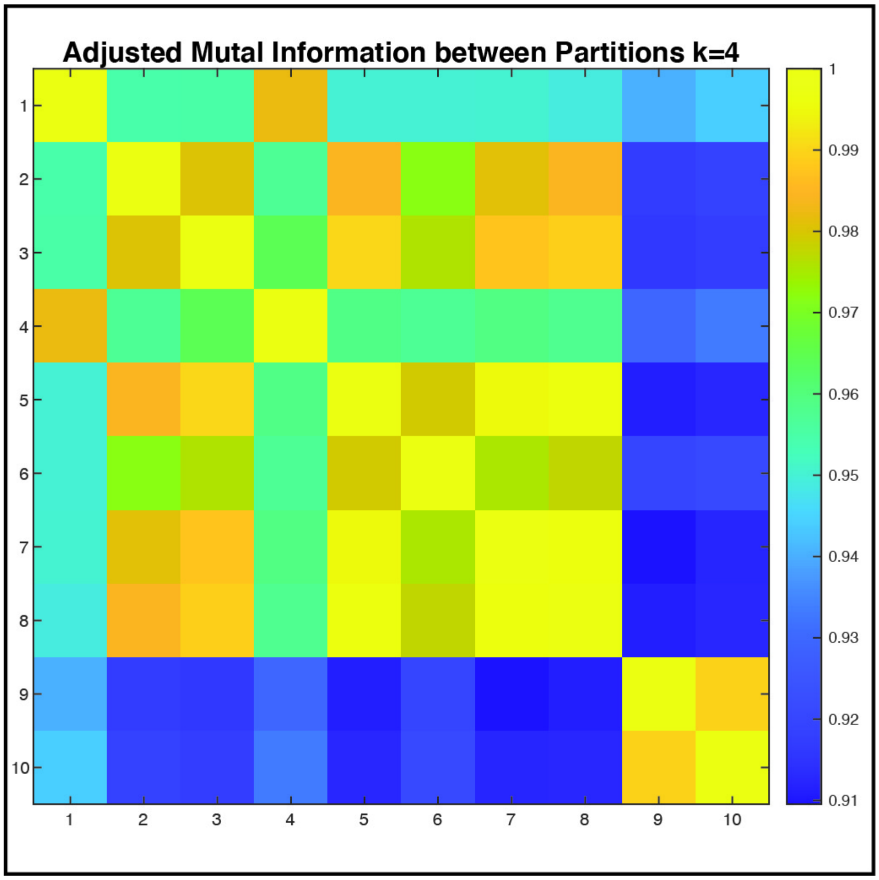


**SI Figure 1: The adjusted mutual information shared between 10 independently generated partitions of our data at *k*=4. Values can range from 0 to 1, with 1 indicating identical partitions**

***Data are clustered in regional activation space beyond null model***

Silhouette scores range from -1 to 1 and are computed for each data point, with 1 indicating that a data point is closer to members of its assigned cluster than to members of the next closest cluster, 0 indicating equidistance between the assigned cluster and the closest cluster, and -1 indicating that the data point is closer to another cluster. In order to show that our data is clustered in regional activation space, we compare silhouette values from our real, k-means clustered data, with those of random Gaussian distributed data, and data from an independent phase randomized (IPR) null model[2] (preserving regional autocorrelation). A two-sample t-test confirms that silhouette values are larger in the real data, indicating that the data are clustered in regional activation space beyond what is expected from a signal with the same regional autocorrelations[3].

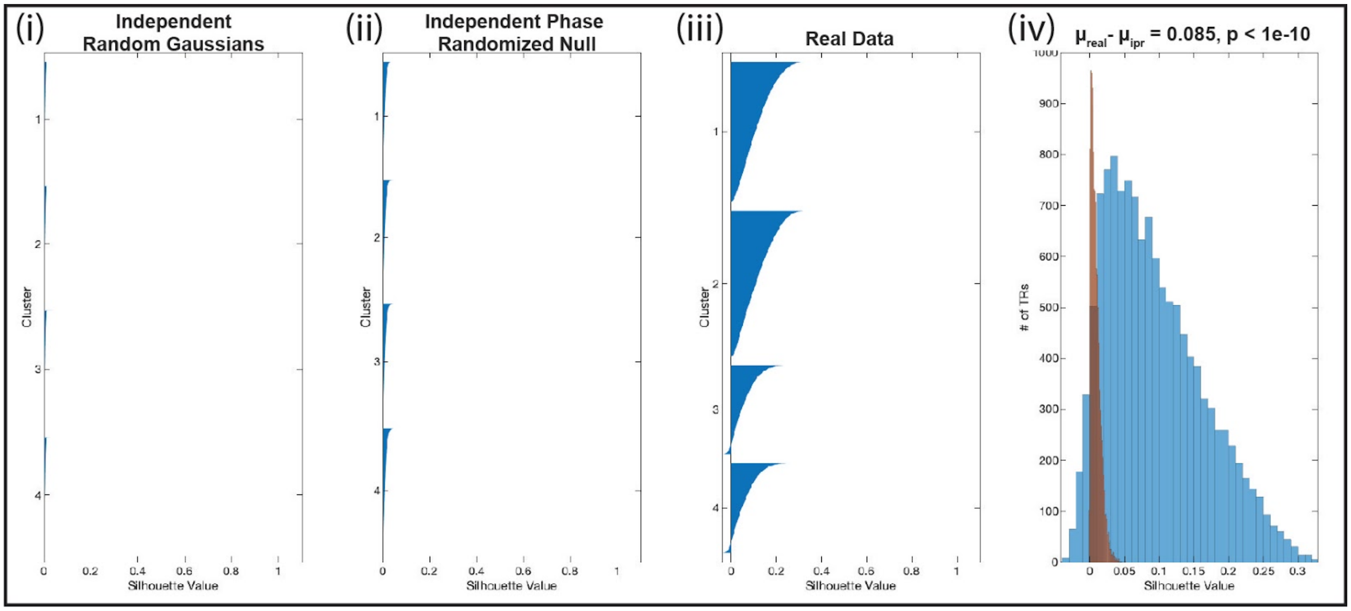


**SI Figure 2:** (i) Silhouette values for data points from 462 independent, normally distributed channels. (ii) Silhouette values for data from an independent phase randomized (IPR) null model applied separately to LSD and placebo BOLD data. This null model preserves regional autocorrelation while eliminating non-stationarities and reducing covariance. (iii) Silhouette values for actual LSD and placebo BOLD data. (iv) Distribution of silhouette values across all clusters for a null model preserving autocorrelation (red, $\mu_{ipr}$) and for actual data (blue, $\mu_{real}$).

***Brain states derived from k-means clustering show 'split' dichotomy***

The brain states used and commented upon in the main text were sometimes referred to as a pair of two 'meta-states', each containing two anticorrelated sub-states. We found that the sub-states were often modified in unison with their partners by LSD. Indeed, this relationship has been previously observed in this type of analysis and it has been observed that two sub-states (e.g. MS-1a and MS-1b) likely correspond to positive and negative spatial principal components of the data[3,4]. This approach has advantages over PCA including the ability to use network control theory. Although their opposite activation patterns can be seen easily on the radial plots and brain volumes in Figure 2, here we show their Pearson correlation values with each other to further illustrate this organization (SI Figure 2, ii).

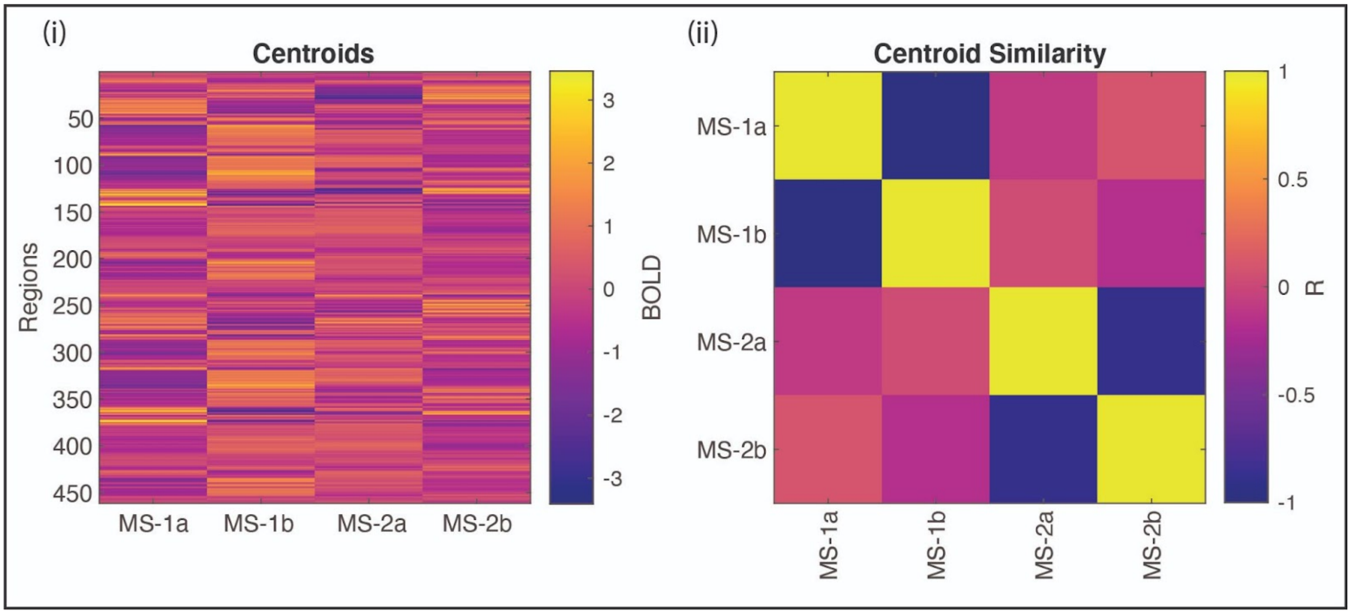


**SI Figure 3: Natural grouping of meta-states based on Pearson correlation.** (i) Centroids of each cluster derived as the mean of all time-points assigned to that cluster. (ii) Pearson correlation values between each cluster. The dark blue squares show the high-values of negative correlations between clusters belonging to the same 'meta-state'.

### ***Psilocybin centroids and correlation results***

Given the relatively small amount of data (9 subjects, 200 frames minutes each) compared to the LSD dataset (15 subjects, 868 frames each), we initialized the k-means clustering algorithm using the group-average centroids derived from the data in the main analysis (Figure 2). We find similar solutions for brain states, and consistent trends and significant findings. Most notably, the energy landscape is significantly lowered in all but one of 16 possible state-transitions (SI Figure 4b, iii). Here, we include the centroids (a), their correlation with the centroids derived from the LSD dataset (b), and the correlations (c) that did not meet significance and thus were not included in Figure 6.

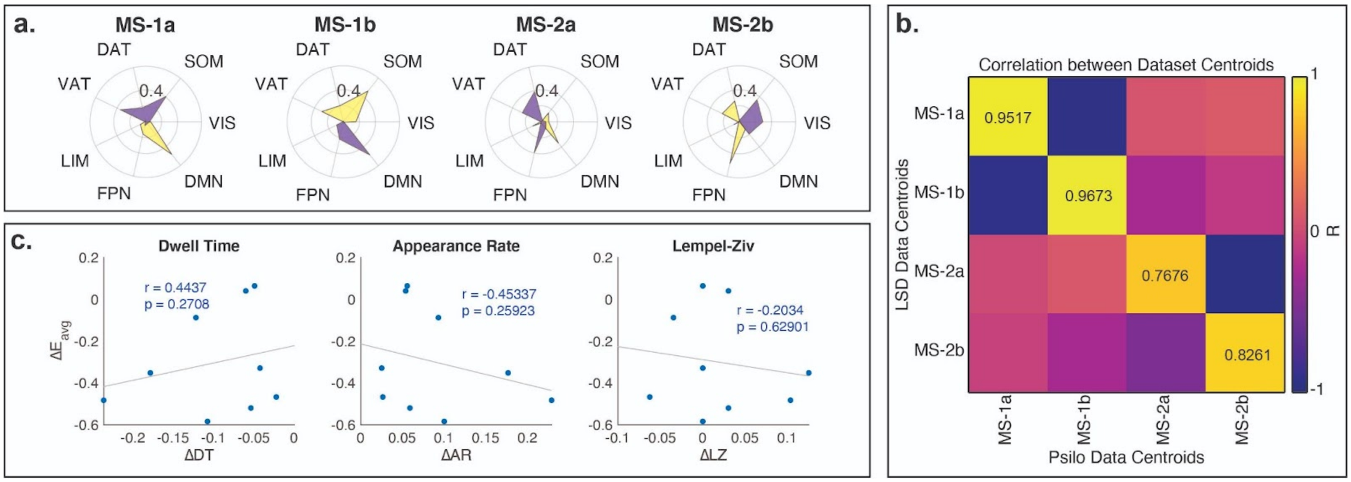


**SI Figure 4: Additional results for the psilocybin data. (a)** Cluster centroids (brain states) derived from the psilocybin data. **(b)** Correlation between the centroids derived from the LSD dataset and the psilocybin dataset (diagonal). **(c)** Pearson correlations between an energy reduction by psilocybin and dynamic measures of dwell time, appearance rate, and Lempel-Ziv. p-values uncorrected.

***Group-level comparison of overall dwell-times and appearance rates for each condition***

When averaging the dwell time of each condition over the four states for each individual, we observe no significant changes between LSD and placebo, even though individual changes in these measures correlated with the amount of energy reduction by LSD. Interestingly, psilocybin did see a significant reduction in average dwell-time and increase in average appearance rate which is easier to interpret under the REBUS hypothesis where decreased priors leads to a flatten landscape and increased cycling between states.

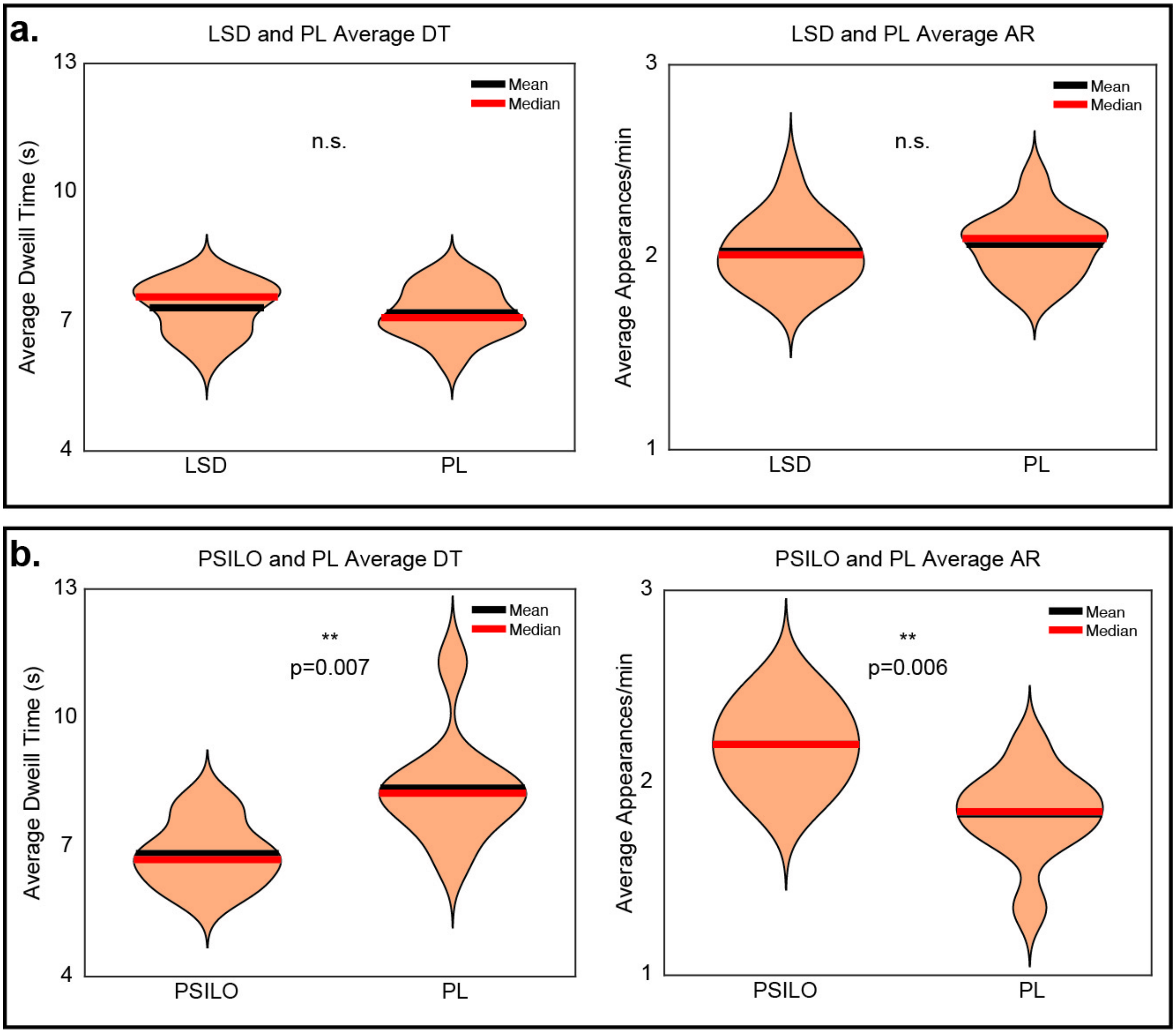


**SI Figure 5: Group-level comparisons of average dwell-times and appearance rates for the (a) LSD and (b) psilocybin datasets.** ** corrected p-values

### ***Comparison of LSD brain state composition to placebo***

Here, we compared the brain states derived from LSD and placebo, finding high consistency between the conditions (mean squared error (MSE) of 0.1141). While the Pearson correlation between the two groups of brain states is high (SI Figure 6d, diagonal), there are some differences (measured in terms of cosine similarity to RSN's) in the states derived from each group. MS-1a has an increase in the high-amplitude activity of the limbic (LIM) network, while MS-1b has an increase in the low-amplitude activity of the LIM network under LSD (SI Figure 6c), together indicating an increase in limbic (bottom-up) representation in the LSD condition. In MS-2 there was a significant increase in the DMN activity in each sub-state (SI Figure 6c). Interestingly, however, LSD increased the amount of DMN activity of amplitude opposite of the DMN's dominant amplitude in each state. In other words, the DMN becomes more polarized (i.e. high and low-amplitude activation levels become further apart).

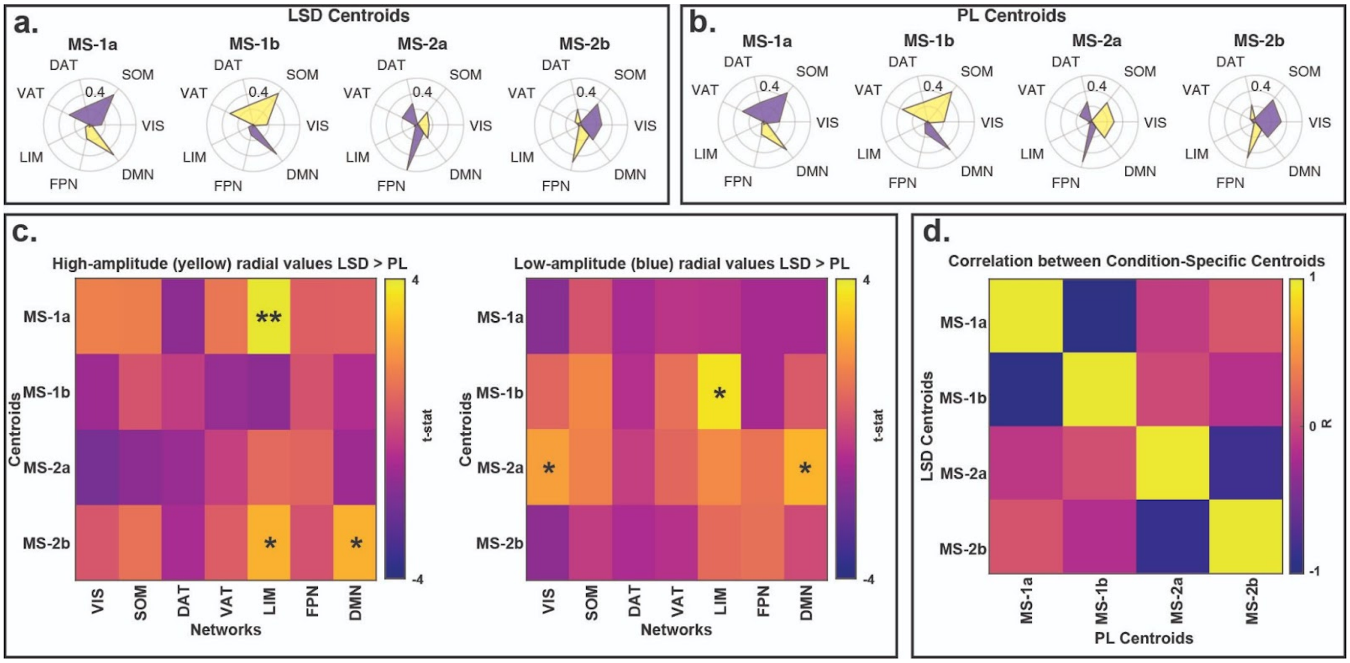


**SI Figure 6: Comparison of condition-specific centroids.** Cosine similarity to RSNs of centroids generated as the mean activation patterns for each cluster **(a)** in the LSD condition and **(b)** in the placebo condition. **(c)** Group level differences in the cosine similarity between LSD and placebo. The only difference that survived correction was a slight increase in the amount of high-amplitude limbic network activation in the MS-1a brain state. This is particularly interesting as it may be representative of the breakdown of the functional hierarchy under psychedelics proposed by REBUS. **(d)** Correlation matrix between the centroids of both conditions shows high agreement between the two conditions (diagonal). The mean squared error between the 4 pairs of states was 0.1141. *significant before multiple comparisons correction, ** significant after multiple comparisons correction.

### ***Within-subject replication of landscape results with music-listening during LSD and placebo administration***

We report our main analysis using resting-state data that did not include exogenous stimulation. However, the same subjects were also scanned while listening to music. Here, we report a replication of our trends with this small subset of the data. Three subjects had technical difficulties with the music, and so n=12 for the following results. For full details on the music condition, see the original publication[5].

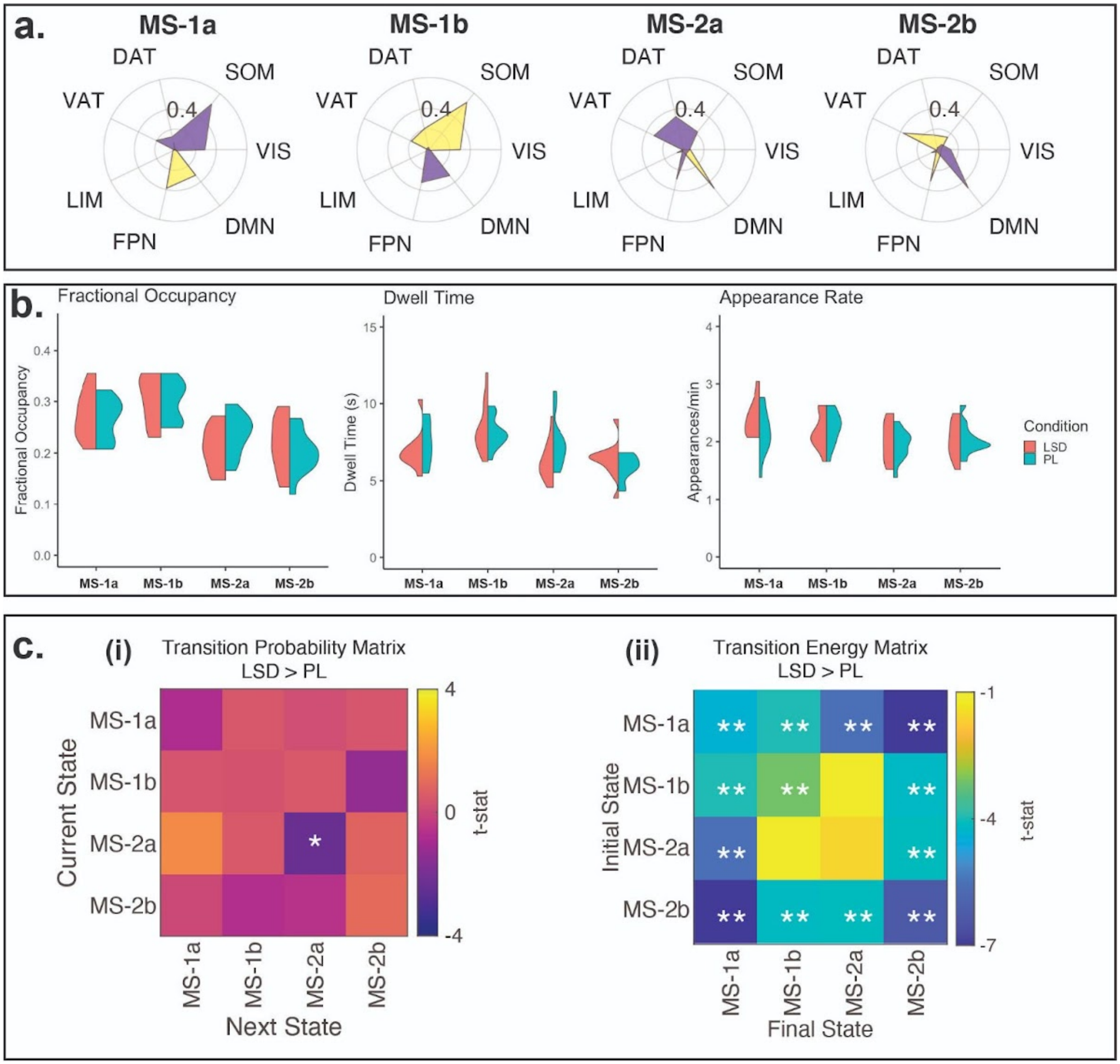


**SI Figure 7: A within-subject replication of the main results using fMRI scans taken while the individuals listened to music.** We find similar brain states to the main resting-state analysis and a flattened energy landscape under LSD. *significant before multiple comparisons correction, ** significant after multiple comparisons correction.

***<u>The effects of parcellation choice, scale, and pre-processing</u>***

To ensure that our results were not specific to our choice of atlas scale, type (anatomical vs functional), and data-processing approach, we replicated the analysis using an alternative parcellation to define the nodes of our brain networks, and using z-scored timeseries prior to clustering. Specifically, we adopted the augmented Schaefer-232 atlas, which is a functional rather than anatomical atlas (obtained by combining the functional local-global cortical parcellation of Schaefer[6] with 200 ROIs and the functional subcortical parcellation of Tian[7]with 32 ROIs); this atlas has been shown to produce brain networks with the most representative topology, and it has been used in recent studies of pharmacologically-induced alterations of brain dynamics[8,9]. SI Figures 8 and 9 show that even with these changes in our analysis workflow, the majority of our results are consistent. The only result that did not hold was the significant correlation between energy reduction and changes in Lempel-Ziv complexity of the brain state sequence, though the direction is still the same. We attribute this to the fact that with this alternative set of analytic choices, there was a group-level increase in direct transitions between MS-1a and MS-1b (SI Figure 9c, i); since these two states are considered aspects of the same meta-state, the metastate-based transformation into a binary sequence required for the calculation of Lempel-Ziv complexity (see *Materials and Methods: Lempel-Ziv Complexity*) would result in fewer observed transitions - thereby making the LSD condition appear less diverse. Importantly, the correlations with changes in state dwell times and appearance rates still hold (SI Figure 10), which is reflective of energy reduction by LSD leading to a more dynamic state-space with shorter dwell times and higher rates of switching.

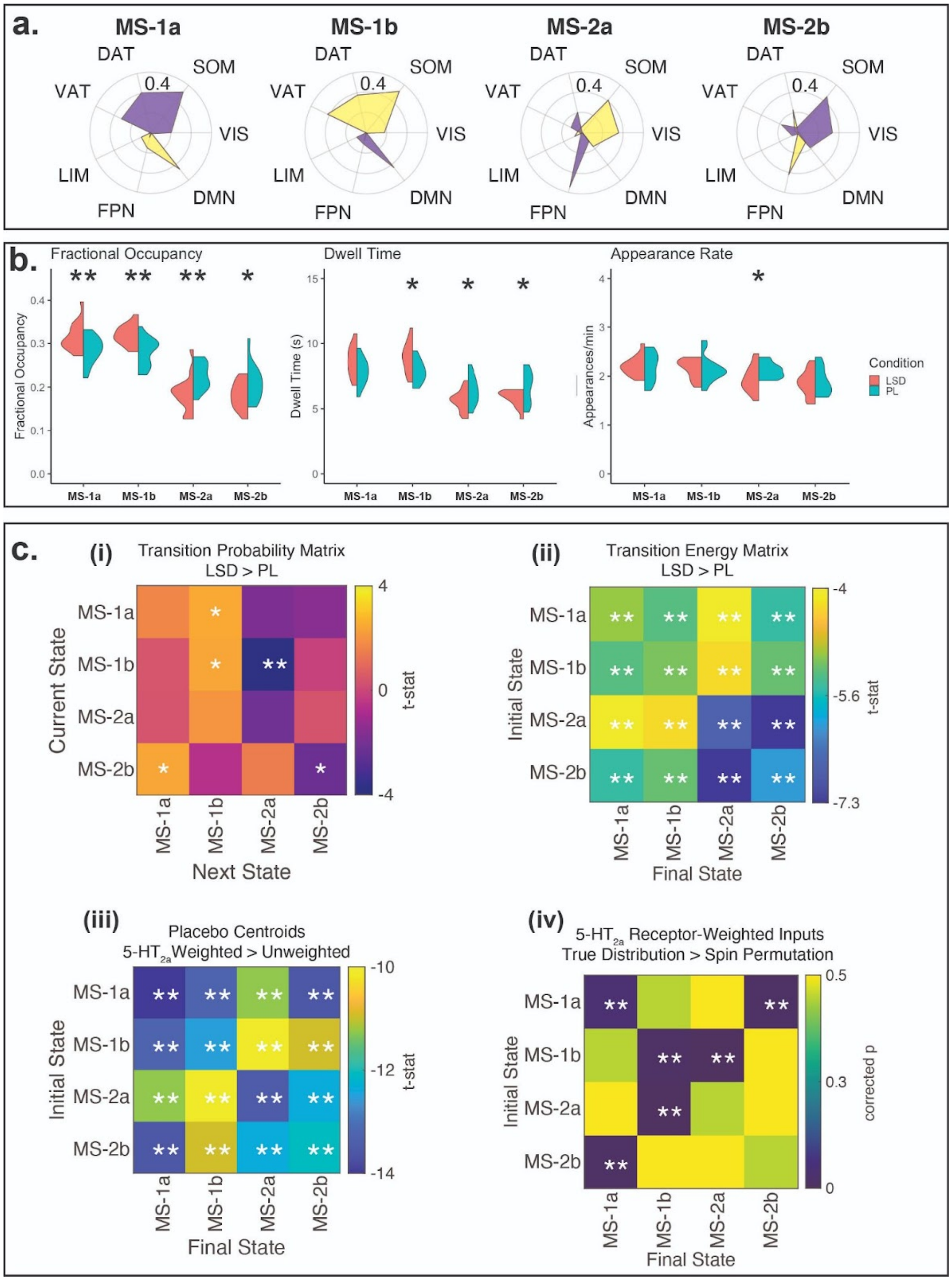


**SI Figure 8: Replication of main results with the augmented Schaefer-232 functional parcellation.**
*significant before multiple comparisons correction, ** significant after multiple comparisons correction.

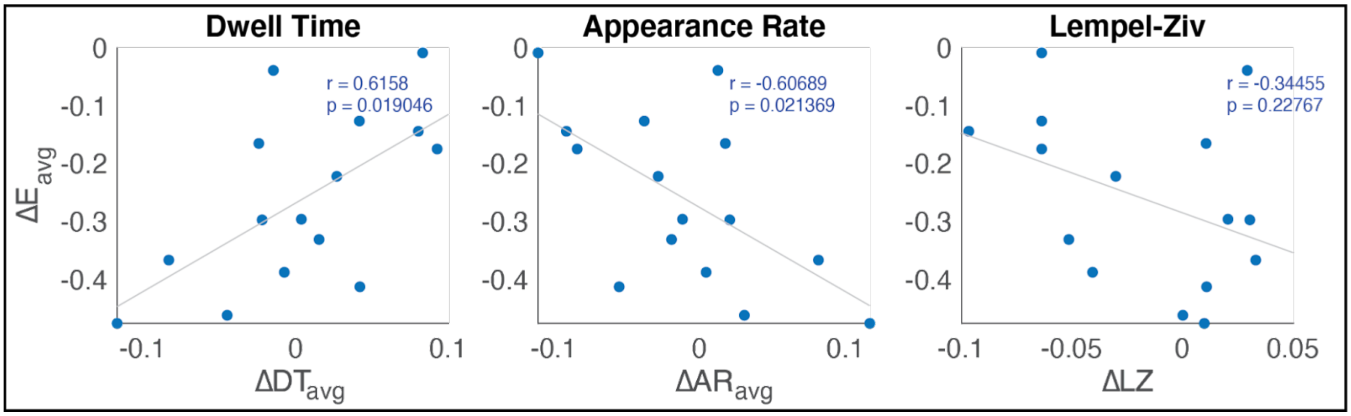


**SI Figure 9: Correlations between energy reduction and changes in state dwell times and appearance rates, as well as the entropy rate, with the augmented Schaefer-232 parcellation.**

The analysis in the main text was performed after de-meaning the regional time-series for each scan, which is not the same as performing global signal regression but it does remove regional and individual level differences in baseline activity. We additionally sought to confirm that global signal regression (GSR) did not alter our main findings (SI Figures 10 and 11).

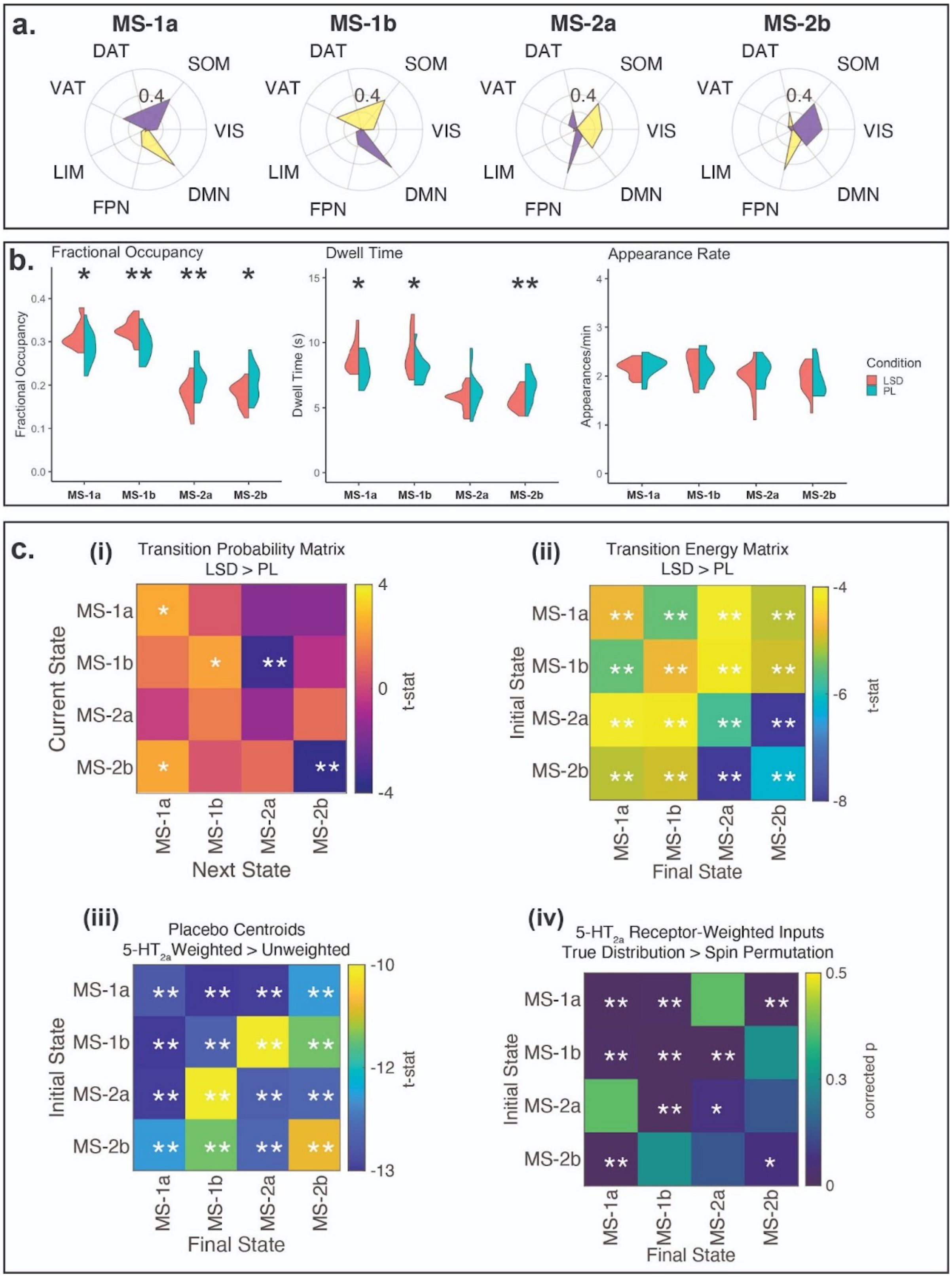


**SI Figure 10: Replication of the main analysis using fMRI data after global signal regression.** *significant before multiple comparisons correction, ** significant after multiple comparisons correction.

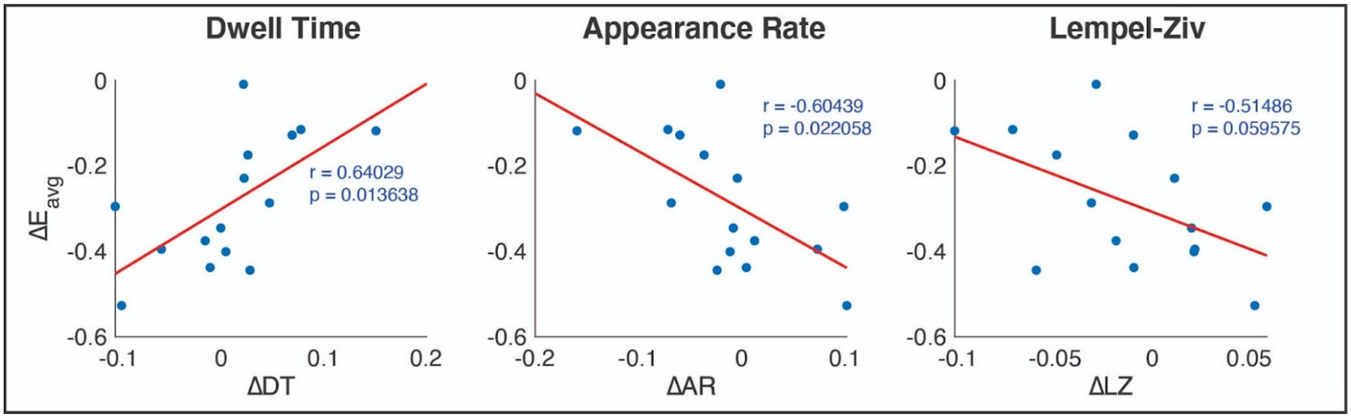


**SI Figure 11: Correlations between energy reduction and changes in state dwell times and appearance rates, as well as the entropy rate, using fMRI data after global signal regression.**

***Brain states derived from Hierarchical Clustering Analysis demonstrate a flattening of the energy landscape***

To ensure that our main findings were not sensitive to the choice of clustering algorithm, we used Hierarchical Clustering Analysis (HCA) with a threshold of four clusters. This produced states comparable to those derived from k-means with k set to 4 (SI Figure 12, a), and also resulted in a flatter energy landscape under LSD compared with placebo (SI Figure 12, b-c).

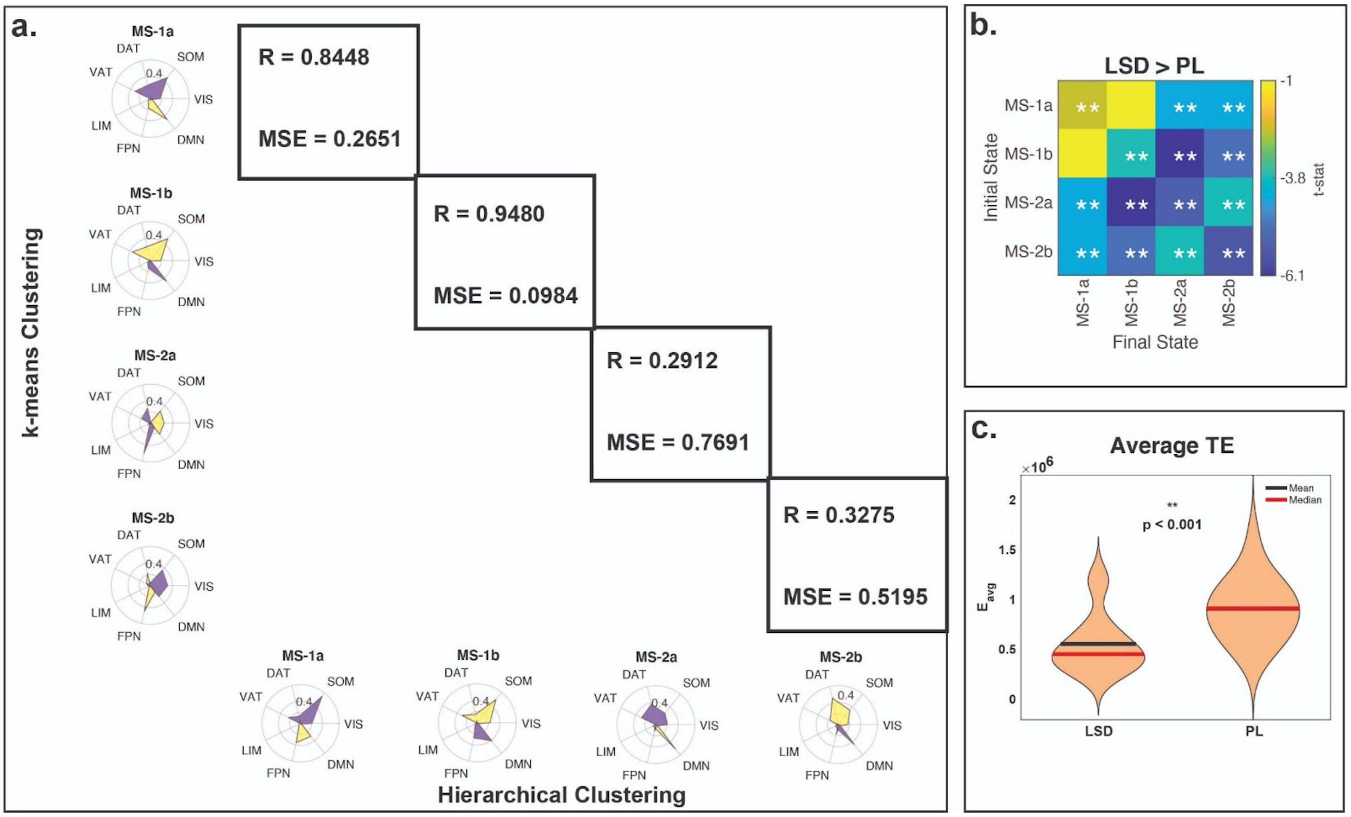


**SI Figure 12: Brain states derived from HCA demonstrate a flattened landscape.** (a) Pearson correlation values and MSE between the 4 states derived from k-means clustering used in the main analysis, and those derived from HCA. (b) Comparison of LSD and placebo energy landscape using the HCA clusters. (c) Comparison of each subject's mean LSD transition energy to their mean placebo transition energy - quantified in each individual as the average energy required to transition between all pairs of states. ** significant after multiple comparisons correction.

***Choosing k***

We performed 50 repetitions of *k*-means clustering for *k*=2 to *k*=14. We quantified the variance explained by clustering as the ratio of between-cluster variance to total variance in the data[3,4,10]. We chose *k* by plotting the variance gained by increasing *k* (SI Figure 13, ii), to observe where increasing *k* begins to provide diminishing returns in terms of variance explained. As has been observed previously[3,4], the elbow of the variance explained curve is in the area of *k*=4-6 (SI Figure 13, i) . In our case specifically, we see that the gain in variance explained by increasing *k* continues to decline following *k*=4 and after *k*=5, the variance gained by increasing *k* falls below 1%, the threshold used previously for choosing *k* (SI Figure 13, ii)[3]. We therefore analyzed the results with *k*=4 and 5 and chose to include *k*=4 in the main analysis for ease of interpretation. We also show that the energy landscape was lowered by LSD for every choice of k (SI Figure 14) and additionally provide the full results for *k*=5 (SI Figures 18 and 19).

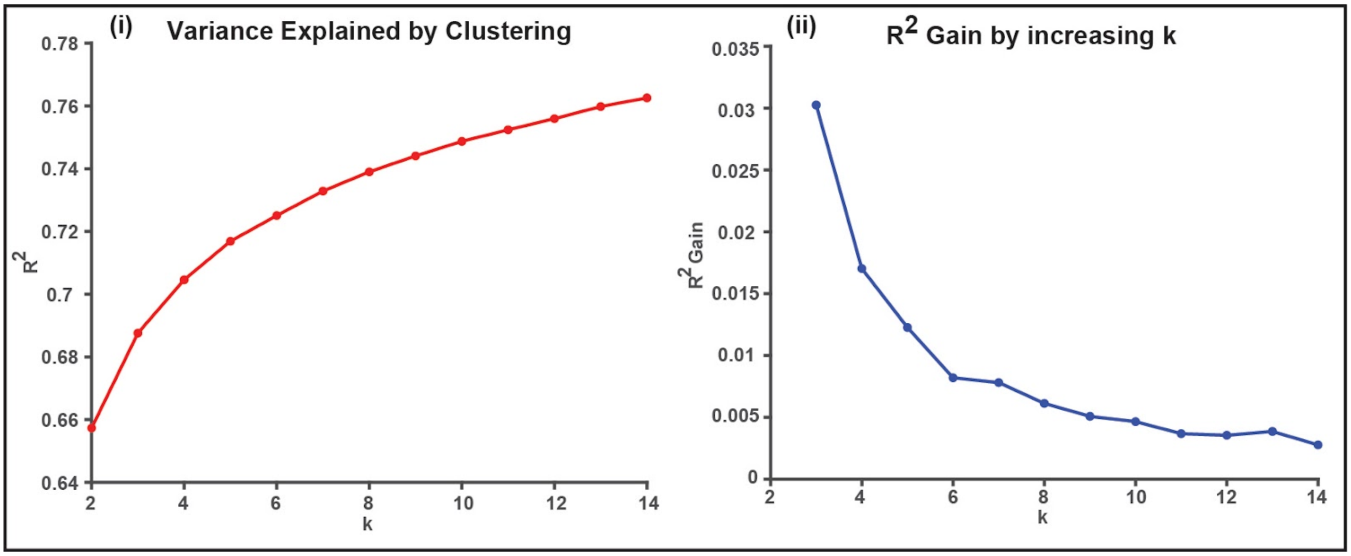


**SI Figure 13: Choosing *k*.** (i) Elbow plot of the variance explained by clustering for each choice of *k*, showing an 'elbow' around 4-6. (ii) Plot of the gained variance explained by increasing *k*. Increasing *k* beyond *k*=5 results in less than a 1% increase of variance explained.

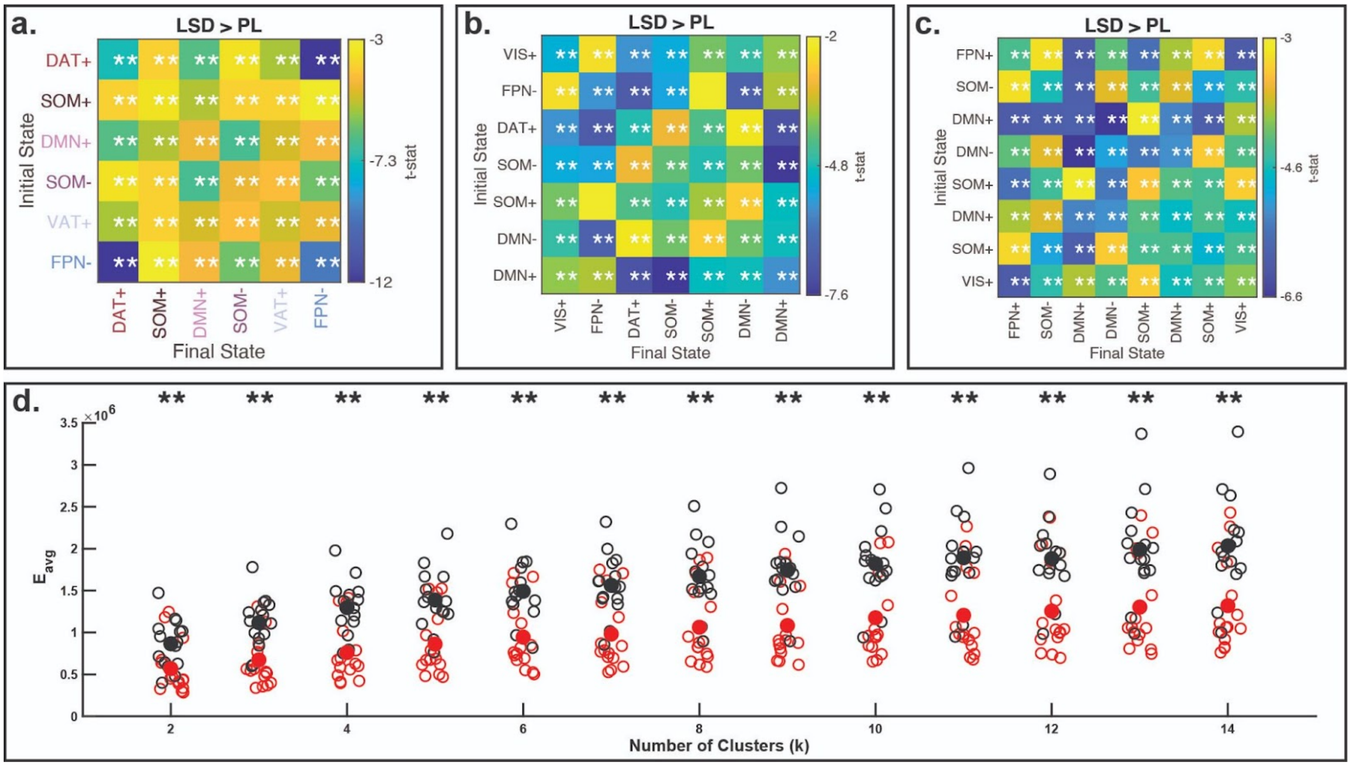

**SI Figure 14: (a-c) Full transition energy comparisons for k=6, 7, and 8, respectively. (d) Average placebo (black) and LSD (red) transition energies for each subject, for k=2:14.** States in a-c are labeled based on their maximum cosine similarity value with the 7 RSN's presented in the radar plots of the main analysis. **(**Filled, condition mean; ** significant after multiple comparisons correction).

***Control energy results are not due to individual or condition-level differences in state representation***

In order to ensure that our results were not an artifact of clustering two dissimilar physiological conditions, we ran k-means on PL and LSD data separately. We matched the resulting centroids derived from each condition based on their maximum correlation with each other (SI Figure 15, i), and then recomputed the transition energy matrix. We replicated the main paper's finding that most transition energies are significantly lower under LSD compared to placebo. In addition, we compared the average transitions energies (over all transitions) between the two conditions and found LSD had significantly lower energies (SI Figure 15, ii). Further, to confirm that differences in control energies are not driven by differences in how well the states represent an individual, we applied k-means to each individual person separately and compared control energies between their respective states. We again ordered centroids based on their correlation to the original 4 clusters in the main analysis and compared individual transitions (SI Figure 15, iii). Again, our main findings hold - LSD lowered the transition energy in all cases. In addition, comparing the average value of the full matrix for the LSD and PL condition we see that overall transition energies are lowered (SI Figure 15, iv).

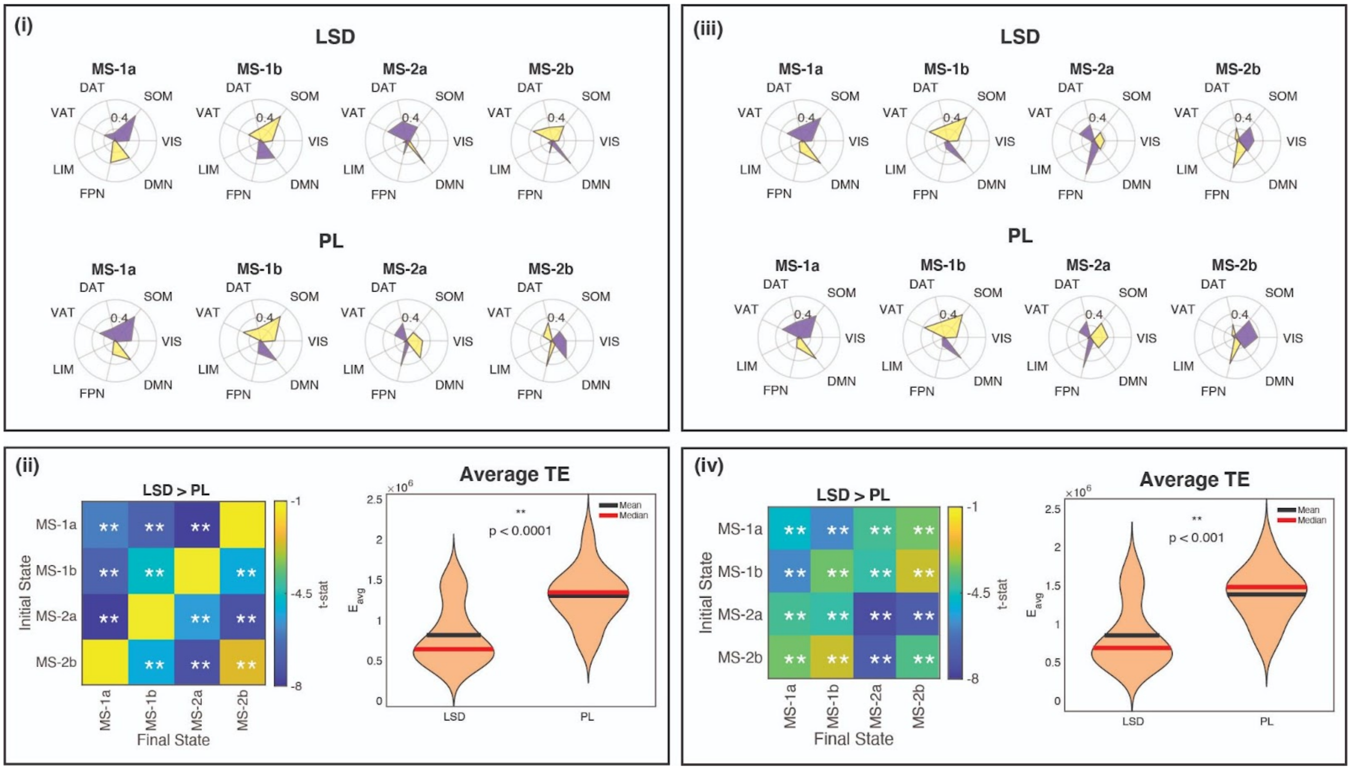


**SI Figure 15: Centroids and energy results from clustering each condition separately (i-ii) and individuals separately (iii-iv).** ** significant after multiple comparisons correction.

### ***Condition-level matrices for results in Figure 5a***

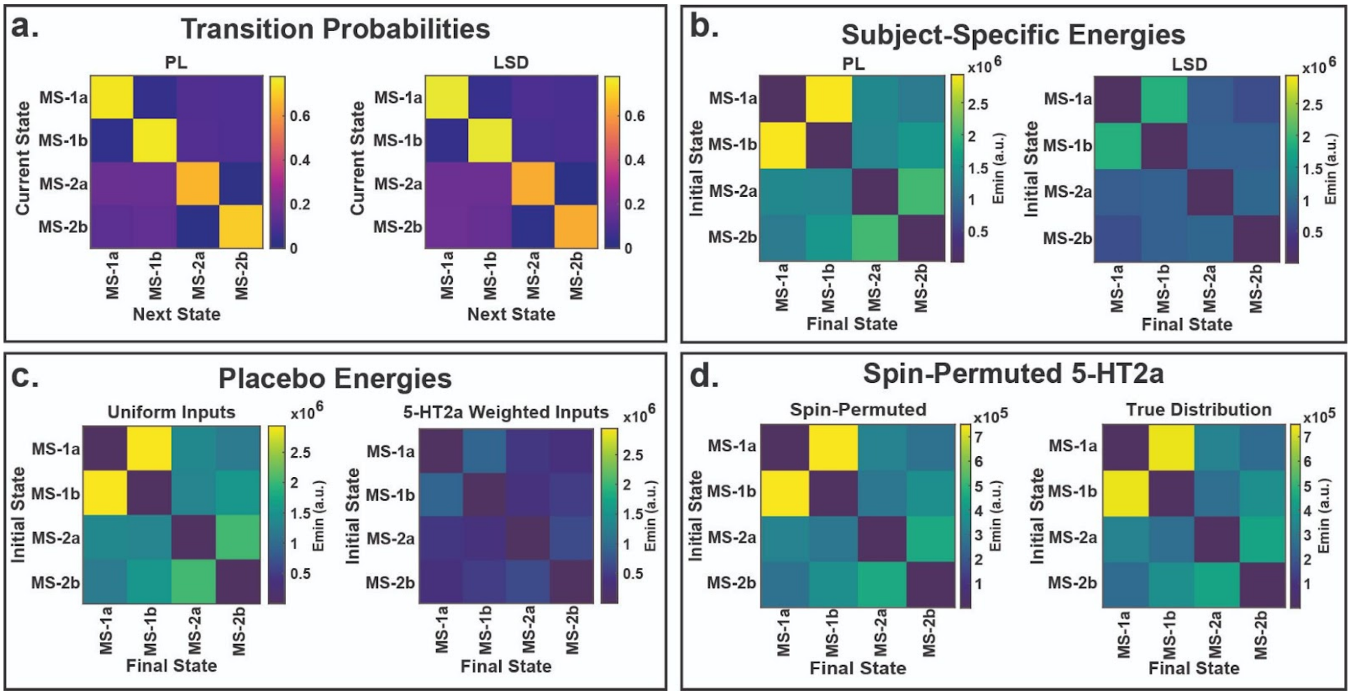


**SI Figure 16: Average group matrices that were used for statistical comparisons in Figure 4a of the main**

**text.**

***Calculating the minimum energy required to transition between states***

For a full discussion on network control theory in the current context we refer the reader to the literature[11–15], and in particular, a recent example by Cornblath et. al[3]. We discuss briefly here the process of choosing *T* - the time over which transitions are allowed - and its implications.

Restating from the main text, we obtained a representative NxN structural connectome *A* obtained using deterministic tractography from HCP subjects (*see Methods and Materials; Structural Connectivity Network Construction*), where N is the number of regions in our atlas. We then employ a linear time-invariant model:

$$\dot{x}(t) = Ax(t) + Bu(t)$$

where x is a vector of length N containing the regional activity at time *t*. *B* is an NxN matrix that contains the control input weights.

To compute the minimum control energy required to drive the system (network) from an initial activity pattern ($x_o$) to a final activity pattern ($x_f$) over some time (*T*), we compute an invertible controllability Gramian (*W*) for controlling the network *A* from the set of network nodes *K* (in our case, every node in the network), where:

$$W = \int_{o}^{T} e^{A(T-\tau)} B B^{\top} e^{A^{\top}(T-\tau)} d\tau$$

where *T* is the time horizon that specifies the time over which input to the system is allowed. After computing the controllability Gramian, we can solve for the minimum control energy $E_{min}$ by computing the quadratic product between the inverted controllability Gramian and the difference between $e^{AT}x_0$ and $x_T$:

$$E_{min} = (e^{AT} x_o - x_T)^{\top} W^{-1} (e^{AT} x_o - x_T)$$

This quantity was calculated for each pair of initial $x_o$ and final $x_T$ brain states to obtain a *k x k* transition energy (TE) matrix, (SI Figure 16, b-d). Following Cornblath et. al.[3], we chose *T* in a data-driven manner. Specifically, we calculated Spearman rank correlations between condition-average transition probability (TP) matrices (SI Figure 16, a) and transition energy (TE) matrices over a range of *T* values (0.001 to 10, increments of 0.5). We chose our final T to be the value that maximized this correlation, which was 0.001 (SI Figure 17).

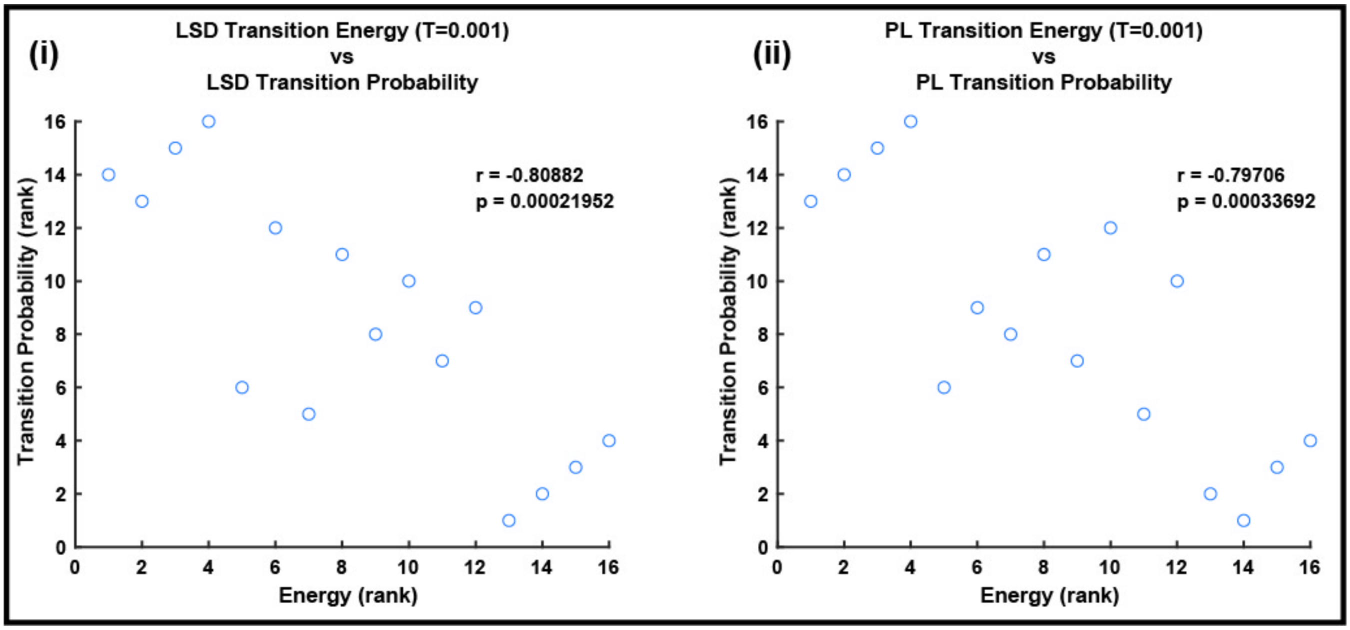


**SI Figure 17: Strongest Spearman correlation between TE and TP occurs at T = 0.001.**

One note: as *T* approaches 0, $E_{min}$ approaches the weighted Euclidean distance between the two states, where the weights are the control inputs in *B,* i.e. $(x_T - x_o)^\top (BB^T)^{-1}(x_T - x_o)$. In this case, the structural connectome becomes less influential on the energy calculations. Previous work has shown generally larger values of optimal *T* than what we found here, but that work was done using task-based fMRI data in controls as well as a different structural connectome derivation. Furthermore, in one of their experiments they did indeed find an optimal *T* of 0.001 as we did in this study, implying weaker influence of structural connectome in navigation of state-space. There could be a number of reasons for our small *T*, not the least of which being that our structural connectome was built from HCP subjects rather than on our individual subjects.

### ***Collection of subjective measures data***

As reported in the original publication[5], VAS (visual analog scale) subjective ratings were obtained after each scan. The scales included items for intensity, simple imagery, complex imagery, positive mood and ego dissolution and emotional arousal. Specifically, they were phrased as follows: 1) “Please rate the intensity of the drug effects during the last scan”, with a bottom anchor of “no effects”, a mid-point anchor of “moderately intense effects” and a top anchor of “extremely intense effects”; 2) “With eyes closed, I saw patterns and colours”, with a bottom anchor of “no more than usual” and a top anchor of “much more than usual”; 3) “With eyes closed, I saw complex visual imagery”, with the same anchors as item 2; 4) “How positive was your mood for the last scan?”, with the same anchors as item 2, plus a mid-point anchor of “somewhat more than usual”; 5) “I experienced a dissolving of my self or ego”, with the same anchors as item 2; and 6) “Please rate your general level of emotional arousal for the last scan”, with a bottom anchor of “not at all emotionally aroused”, a mid-point anchor of “moderately emotionally aroused” and a top anchor of “extremely emotionally aroused”.

At the end of each scanning day, subjects also completed the 11-item Altered States of Consciousness (ASC) scale[16,17], providing retrospective subjective evaluations pertaining to the peak of the experience (i.e. during fMRI scanning).

***Results are replicated with k=5***

Previous work has chosen *k*=5 for their analysis[3]. We performed our analysis with *k*=5 as well to demonstrate that our results are robust. Interestingly, at *k*=5, the ventral attention network (VAT), is isolated as a state. All major results and trends hold (SI Figures 18 and 19). Note that we did not calculate the Lempel-Ziv complexity for this clustering solution, since the only principled way to do so would be to collapse the VAT+ state with the two SOM states, thereby coinciding with the analysis performed for the 4-state clustering solution.

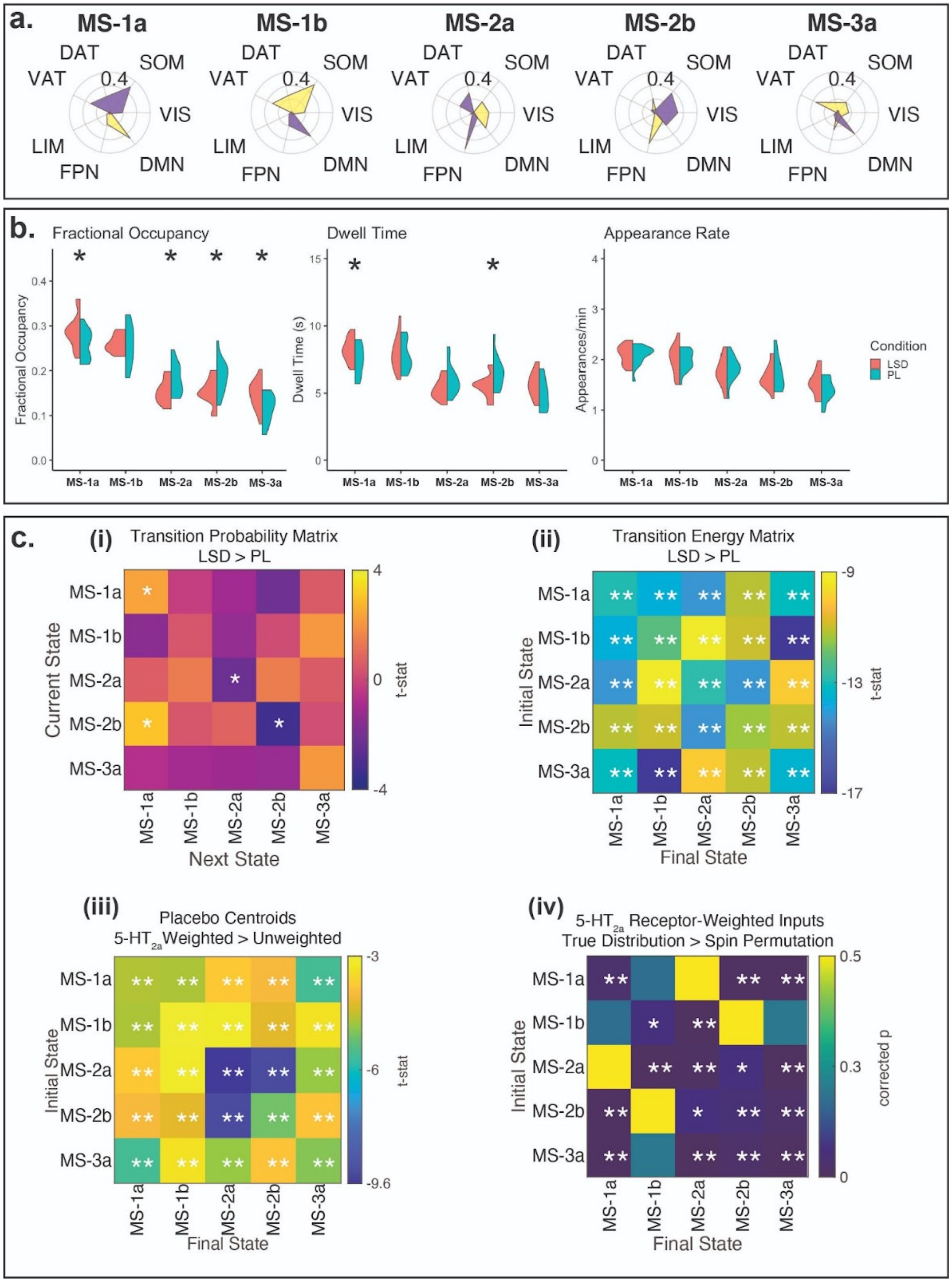


**SI Figure 18: Replication of main analysis with *k*=5.** *significant before multiple comparisons correction, ** significant after multiple comparisons correction.

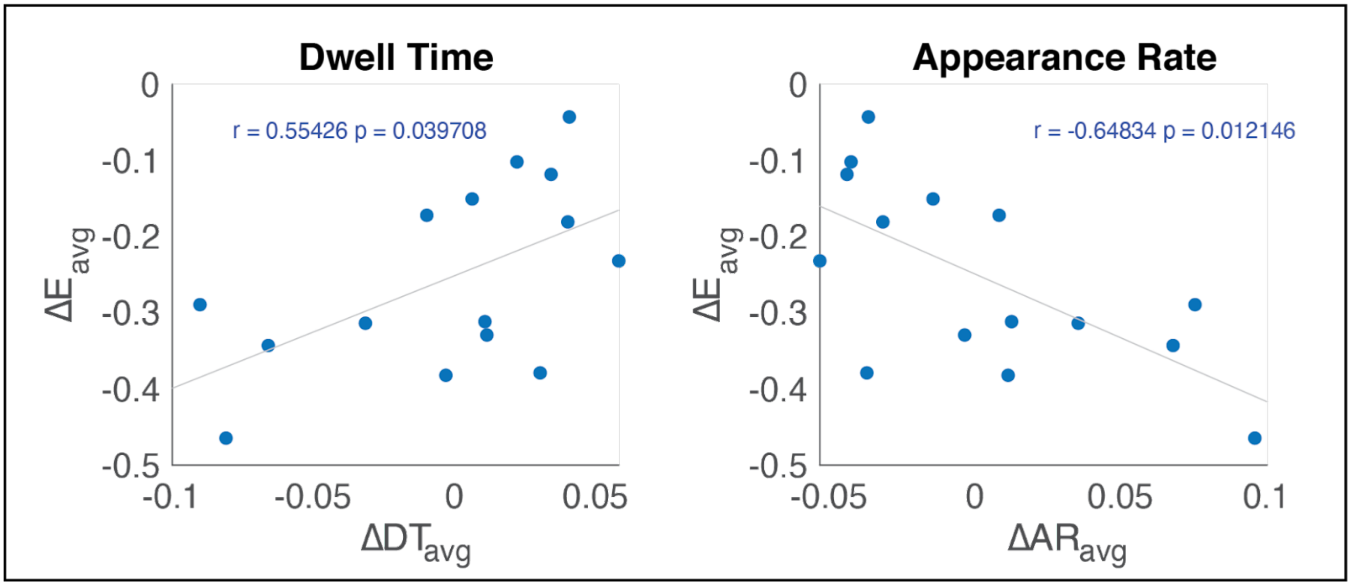


**SI Figure 19: Correlations between energy reduction and changes in state dwell times and appearance rates for *k*=5.**

***Motion***

The motion quality of these data and the measures taken have been detailed extensively in the supplemental information of the original publication[5,18]. In addition to motion correction and scrubbing (FD > 0.4), de-spiking was performed because it has been shown to improve motion-correction and create more accurate FD values[19] and low-pass filtering at 0.08 Hz was performed because it has been shown to perform well in removing high frequency motion[20]. The six motion regressors were used as covariates in linear regression. It was decided that using more than six (e.g., “Friston 24-parameter motion regression”)[21] would be redundant and may impinge on neural signal[22] (especially when other rigorous processes such as scrubbing[23] and local white matter (WM) regression were applied[24]) . Using anatomical regressors is also a common step to clean noise and ventricles, draining vien (DV) and local WM were used in the pipeline employed in the present analyses. Local WM regression has been suggested to perform better than global WM regression[19]. In the main text, our correlations in Figure 6 (and the replication correlations between the same metrics in the SI) were performed using the difference in mean FD between drug and placebo as a covariate of non-interest. We now go a step further and correlate each subject’s mean FD with their fractional occupancy, dwell time, and appearance rate in each cluster, separately for each condition in the LSD dataset (SI Figures 20 and 21). We do not find any significant correlations. Additionally, we do not find any significant correlation between a subject’s mean FD and their average transition energy for either condition (SI Figure 22). We also replicated our main findings after applying GSR, which has been shown to further minimize motion effects since the global signal has been shown to contain signal related to motion (although we note that it also contains neuronal signal of relevance[25]).

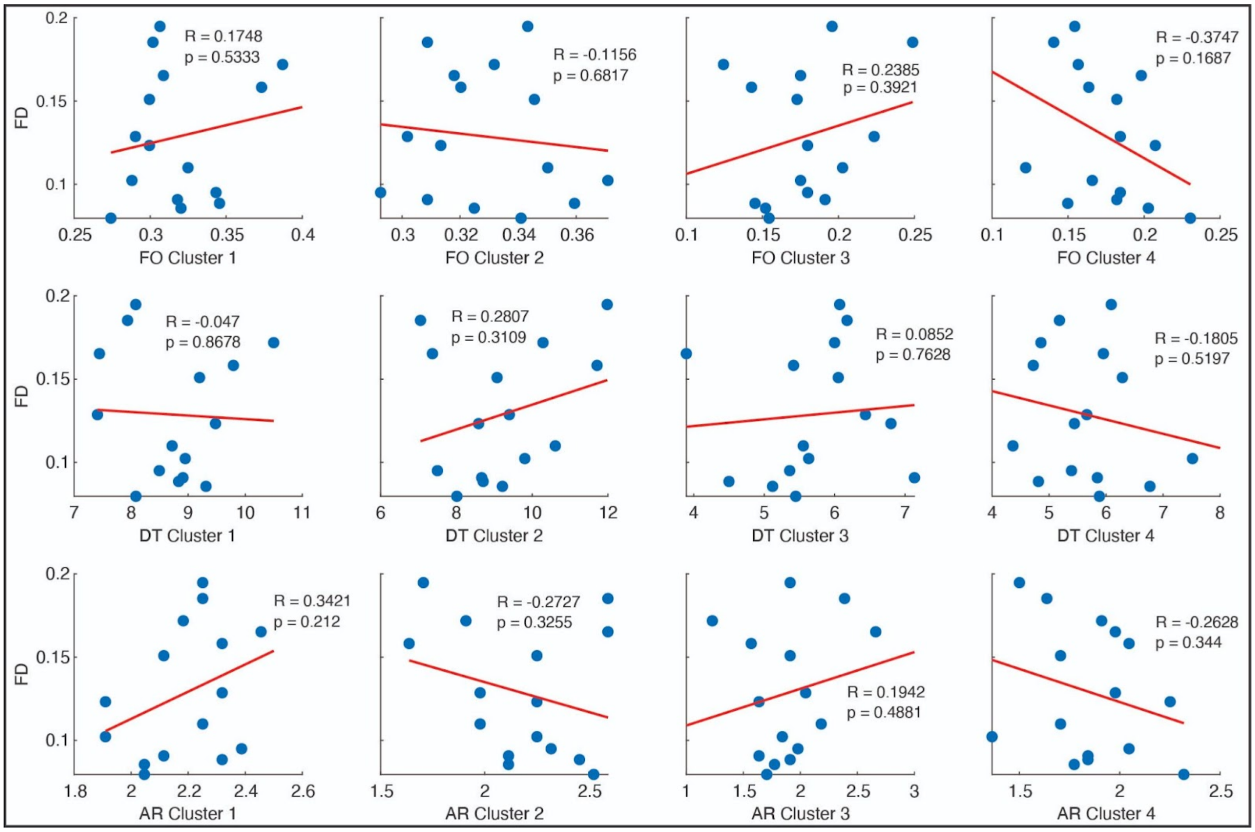


**SI Figure 20: Mean framewise-displacement (FD) of the LSD condition correlated with fractional occupancy (FO), dwell time (DT), and appearance rate (AR).** No significant correlations were found, p-values uncorrected.

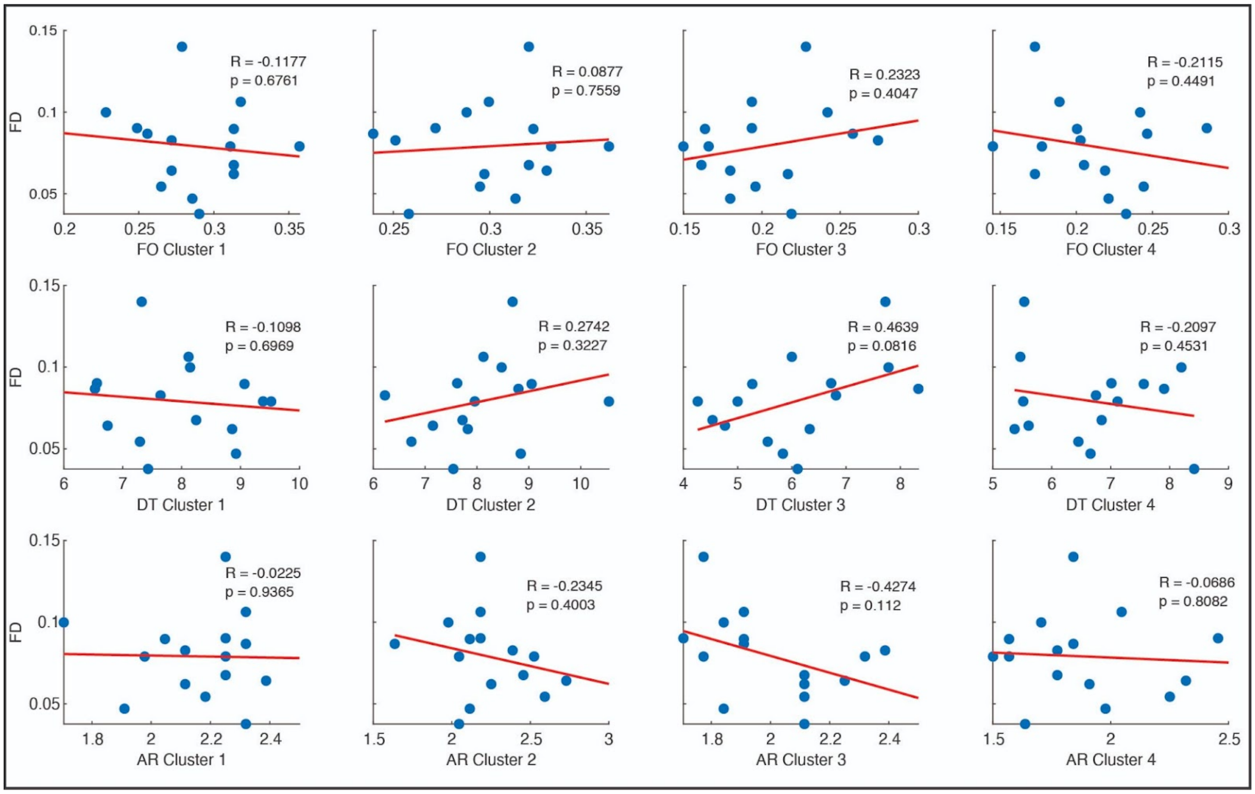


**SI Figure 21: Mean framewise-displacement (FD) of the placebo condition correlated with fractional occupancy (FO), dwell time (DT), and appearance rate (AR).** No significant correlations were found, p-values uncorrected.

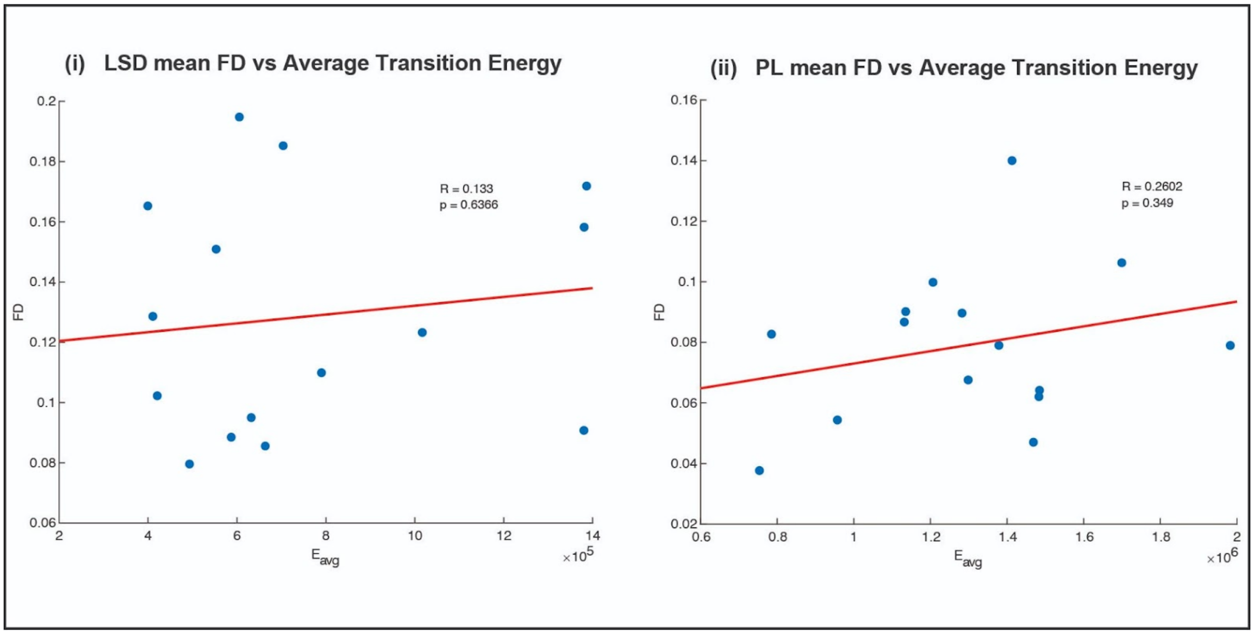


**SI Figure 22: Mean framewise-displacement of the LSD (i) and placebo (ii) condition correlated with the average transition energies of each condition.** Neither correlation is significant, p-values uncorrected.

***Transition probability results when excluding self-transitions***

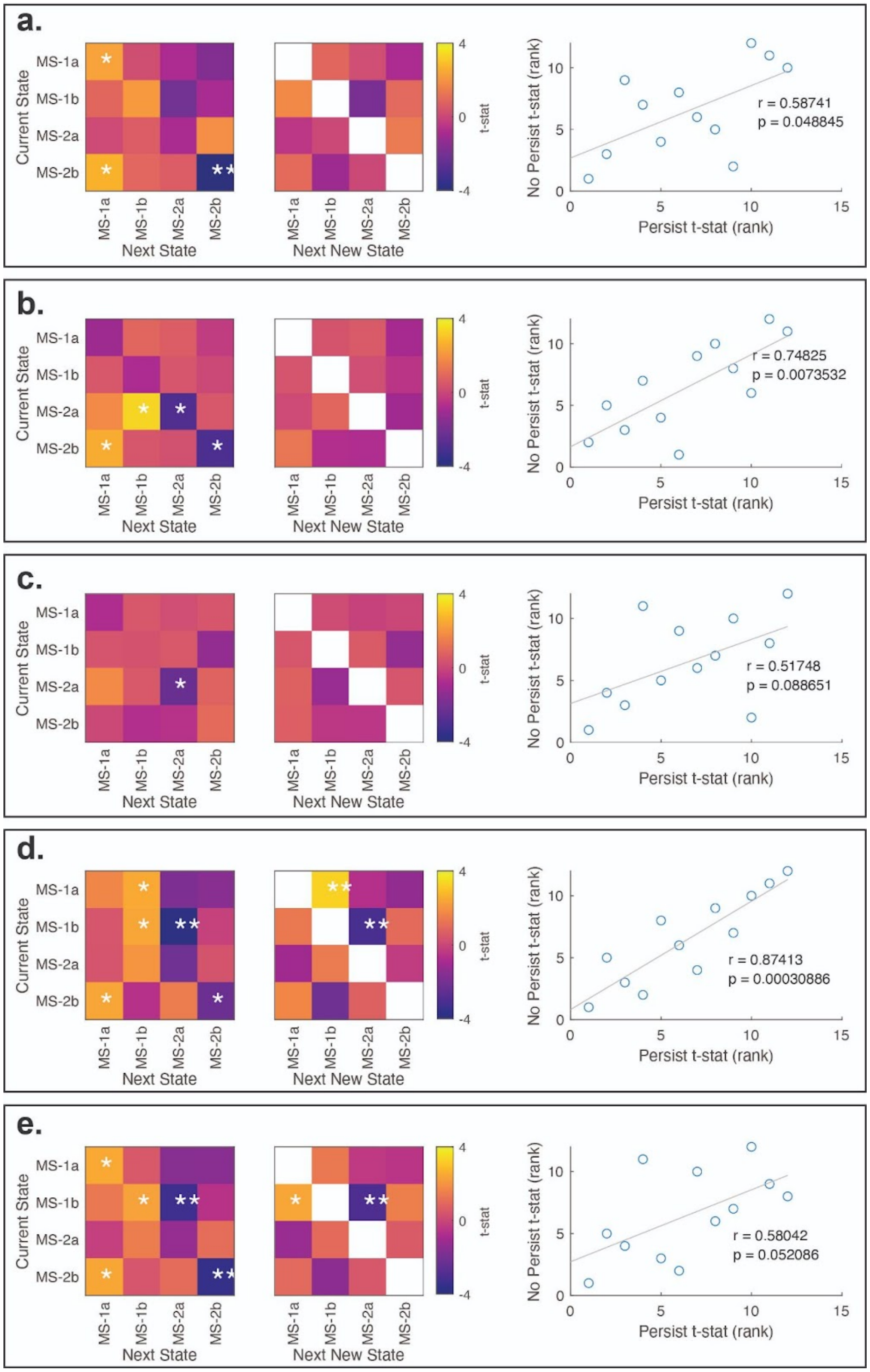


**SI Figure 23:** Our main results and the replications present the transition probabilities that include the probability of staying in the same state (persistence). Here, we present these results (persist; left column) alongside the results when excluding self-transitions (no persist; middle column), and we correlate the off-diagonal of the results obtained from each version (persist vs no persist; right column). **(a)** LSD main results, **(b)** psilocybin main results, **(c)** music replication, **(d)** z-scored sch232 replication, **(e)** GSR replication. *significant before multiple comparisons correction, ** significant after multiple comparisons correction, correlation p-values are uncorrected.
